# A Comprehensive Evaluation of Large Language Models in Mining Gene Interactions and Pathway Knowledge

**DOI:** 10.1101/2024.01.21.576542

**Authors:** Muhammad Azam, Yibo Chen, Micheal Olaolu Arowolo, Haowang Liu, Mihail Popescu, Dong Xu

**Affiliations:** Department of Electrical Engineering and Computer Science, University of Missouri, Columbia, Missouri, USA; Bond Life Sciences Center, University of Missouri, Columbia, Missouri, USA; Institute for Data Science and Informatics, University of Missouri, Columbia, Missouri, USA; Department of Biomedical Informatics, Biostatistics and Medical Epidemiology, University of Missouri, Columbia, Missouri, USA

**Keywords:** Large Language Model, Gene-gene Interaction, KEGG Pathway, Biomedical Text Mining

## Abstract

**Background:** Understanding complex biological pathways, including gene-gene interactions and gene regulatory networks, is critical for exploring disease mechanisms and drug development. Manual literature curation of biological pathways is useful but cannot keep up with the exponential growth of the literature. Large-scale language models (LLMs), notable for their vast parameter sizes and comprehensive training on extensive text corpora, have great potential in automated text mining of biological pathways.

**Method:** This study assesses the effectiveness of 21 LLMs, including both API-based models and open-source models. The evaluation focused on two key aspects: gene regulatory relations (specifically, ‘activation’, ‘inhibition’, and ‘phosphorylation’) and KEGG pathway component recognition. The performance of these models was analyzed using statistical metrics such as precision, recall, F1 scores, and the Jaccard similarity index.

**Results:** Our results indicated a significant disparity in model performance. Among the API-based models, ChatGPT-4 and Claude-Pro showed superior performance, with an F1 score of 0.4448 and 0.4386 for the gene regulatory relation prediction, and a Jaccard similarity index of 0.2778 and 0.2657 for the KEGG pathway prediction, respectively. Open-source models lagged their API-based counterparts, where Falcon-180b-chat and llama1-7b led with the highest performance in gene regulatory relations (F1 of 0.2787 and 0.1923, respectively) and KEGG pathway recognition (Jaccard similarity index of 0.2237 and 0. 2207, respectively).

**Conclusion:** LLMs are valuable in biomedical research, especially in gene network analysis and pathway mapping. However, their effectiveness varies, necessitating careful model selection. This work also provided a case study and insight into using LLMs as knowledge graphs.

## 1. INTRODUCTION

Biological pathways, encompassing gene-gene interactions, metabolic networks, and gene regulatory networks, are complex systems integral to processes like signaling [1]. Their understanding is crucial in deciphering disease mechanisms and advancing drug development. Biological pathway information is contained in the literature. Manual curations of the literature produced databases like Kyoto Encyclopedia of Genes and Genomes (KEGG) [2], which are instrumental in systematically organizing and visualizing these networks. However, extracting knowledge from the biomedical text is labour-intensive and time-consuming, which led to the rise of automated mining techniques in distilling valuable insights from the extensive biomedical literature [4].

A major development in natural language processing is the emergence of large-scale language models (LLMs), characterized by their enormous parameter sizes and training on extensive text corpora [5]. Their ability to generate coherent, contextually relevant text makes them particularly suitable for biomedical text generation and mining [6]. Models like BioLinkBERT, which utilizes a domain-specific T5 model trained on extensive biomedical corpora, have shown the potential of LLMs in biomedical text mining [7]. The “pre-train, prompt, and predict” paradigm is an emerging approach in such an LLM application, involving enhancing problem statements with specific instructions for learning from limited examples in prompts [8]. Among the most recent LLMs, ChatGPT-4 is recognized for its advanced language comprehension and generation abilities, enhancing conversational AI, code generation, and language translation tasks. Its predecessor, ChatGPT-3.5, established the groundwork for these advanced tasks [9]. Claude-2 and Claude Instant have shown proficiency in context-aware responses, especially in extended interactions [10], while the Cohere Playground excels in text classification, sentiment analysis, and summarization [11]. In the open-source LLMs, Codellama-34instruct is known for its efficiency in instruction-based tasks [12], and the WizardLM series, including WizardLM-70b and WizardLM-13b, excel in knowledge retrieval and processing [13]. Falcon-180b-chat specializes in conversational AI [14], and Mistral-7b-instruct is tailored for instruction-following tasks. The Vicuna series, including Vicuna-7b, Vicuna-33b, and Vicuna-13b [15], and the Llama2 series, with models like Llama2-13b, Llama2-7b, and Llama2-70b [16], are used in a variety of language modelling and text generation applications. Qwen-14b is recognized for its efficiency in medium-scale language tasks [17].

Although people often use LLMs as knowledge bases or knowledge graphs to retrieve information, their accuracies are rarely studied, even when ground-truth data are available. It creates uncertain confidence and unclear directions to improve. To address this issue in a case study, we aim to evaluate the capacities of LLMs for extracting crucial biomedical information, such as pathway knowledge, gene interactions, and regulatory details. We conducted a comprehensive evaluation of both API-based models (e.g., ChatGPT-4 and Claude-2) and open-source models (e.g., Codellama-34instruct and Falcon-180b-chat) in their capacities in predicting gene regulatory relationships and KEGG pathway components.

The remainder of this paper is structured as follows: Section 2 presents the results of our experiments. Section 3 summarizes key findings and outlines future research directions. Finally, Section 4 provides method details.

## 2. RESULTS

We conducted a two-part analysis, the first focusing on gene regulatory relations (specifically “activation”, “inhibition”, and “phosphorylation”) and the second on KEGG Pathway Recognition (predicting gene names in the form of strings in a pathway). We critically evaluated 21 LLMs against a dataset from the KEGG pathway, which included 200 genes and their known relationships. Our approach involved using specially crafted prompts to extract data from each LLM. The performance of these models was then rigorously assessed using key statistical indicators such as Precision, recall, and F1 scores. These measures provided a comprehensive understanding of each model’s effectiveness in accurately identifying and interpreting gene regulatory relations. We conducted a similar assessment for KEGG Pathway Recognition, i.e., to retrieve genes involved in a given pathway. The insights garnered from this study offer a detailed perspective on the capabilities and limitations of LLMs in gene regulatory analysis.

### 2.1 Evaluations of Gene Regulatory Relations

Figure 1 assesses the effectiveness of various LLMs for identifying gene regulatory relationships encompassing “activation”, “inhibition”, and “phosphorylation”. A distinct performance gradient was observed between proprietary API-based models and open-source alternatives. Among the API-based LLMs, ChatGPT-4 emerged as the standout performer, exhibiting the highest F1 score of 0.4448 alongside commendable recall and precision rates, both measured at 0.3881 and 0.5307. A close second, Claude-Pro registered a recall and precision, complemented by a of 0.3781 and 0.5305 and an F1 score of 0.4386, underscoring its efficacy in the domain.

**Figure 1:**
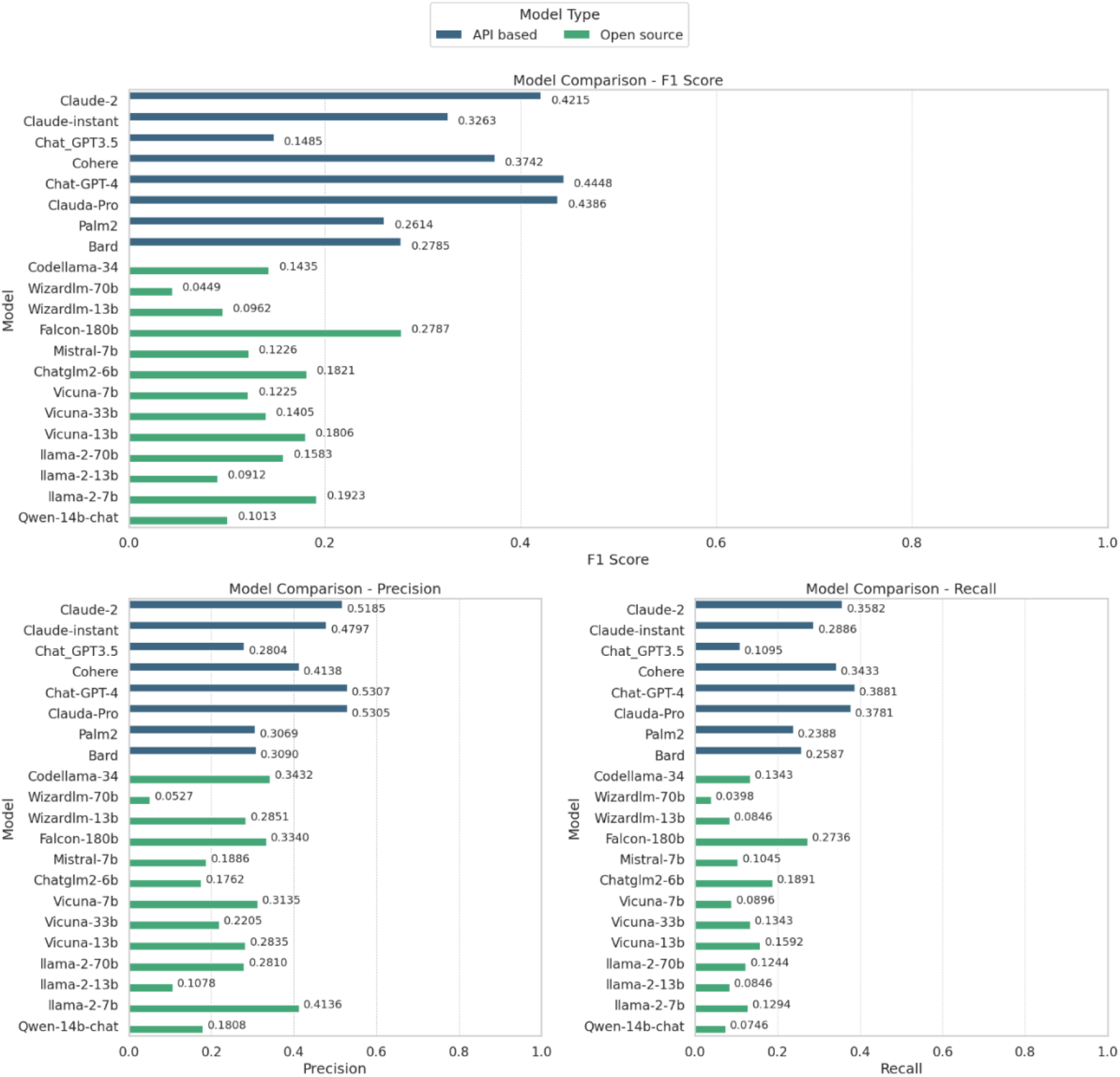
Comparative analysis of gene prediction accuracy across API-based and open-source models, measured by Precision, Recall, and F1 Score.

Conversely, ChatGPT-3.5 displayed considerable limitations in this specific analytical setting, with a recorded Recall of 0.1095 and an F1 score of 0.1485, one of the poorest performers among 21 LLMs. The remainder of the API-based models, including Cohere, Claude-2, Palm2, Bard, and Claude-instant, demonstrated moderate capabilities, with their performance metrics clustering below the 0.4 threshold across all four measured indices. On the open-source front, the results were more varied, with none of the models surpassing the API-based frontrunners. Llama-2-7b highlighted the highest precision among its open-source peers at 0.4136 (i.e., with the least hallucination), although it yielded modest recall and F1 scores, with 0.1923 and 0.129, respectively. This discrepancy points to a considerable number of missed true positives. The rest of the open-source models, including but not limited to Codellama-34, Mistral-7b, vicuna-33b, vicuna-13b, Llama-2-70b and falcon-180b, demonstrated lower levels of recall and precision, with corresponding F1 scores below the 0.30 mark. Models such as wizardly-70 b, wizardly-13b, Vicuna-7b, Llama-2-7b-chat, llama-2-13b, and Qwen-14, occupied the lower tier of performance, with Qwen-14b-chat at the bottom (0.1500 across all categories). API-based models, particularly Claude-Pro and ChatGPT-4, demonstrated superior performance in detecting gene regulatory relationships when compared to the open-source models tested.

### 2.2 Evaluations of KEGG Pathway Recognition

In the second part of our study, we focused on the recognition of seven key biomedical terms within the KEGG Pathway: Adherens junction, Tight junction, GAP junction, Cellular senescence, Phagosome, Proteoglycans in cancer, and Autoimmune thyroid disease. We employed the 21 LLMs to assess their performance in identifying genes associated with these biomedical terms. To refine our analysis, we ran each query five times using the default settings of the 21 LLMs and assigned weights to the genes based on their occurrence. We used the Jaccard similarity index to compare the genes matched by the models with those in the established ground truth.

#### 2.2.1. Adherens Junction complex

Figure 2 shows the assessment of Adherens junction complex, a pivotal factor in cellular adhesion and signal transduction. Figure 2A presents a comparison of the 21 models. The figure maps the prediction accuracies of each model in identifying true genes associated with the Adherens junction, including CDH1, CTNNB1, CTNNA, CTNND1, CDC42, CDH3, ACTN4, ACTB, EP300, FGFR1, LEF1, PARD3, MET, LEF1, and SMAD4. It also quantifies the likelihood for each gene to be predicted correctly within the respective models, which varies greatly. CDH1 was predicted correctly by all models, while some genes (MET, SMAD4, and LEF1) could not be predicted at all by any model. This may reflect the available of relevant text. For example, when we search “CDH1 Adherens” together at Pubmed (https://pubmed.ncbi.nlm.nih.gov) on January 15, 2024, 150 hits were obtained; in contrast, the search of “MET Adherens”, “SMAD4 Adherens” and “LEF1 Adherens” yielded only 50, 10 and 26 hits, respectively (“MET” had more hits probably because it is not a unique name and it also represents amino acid methionine).

**Figure 2:**
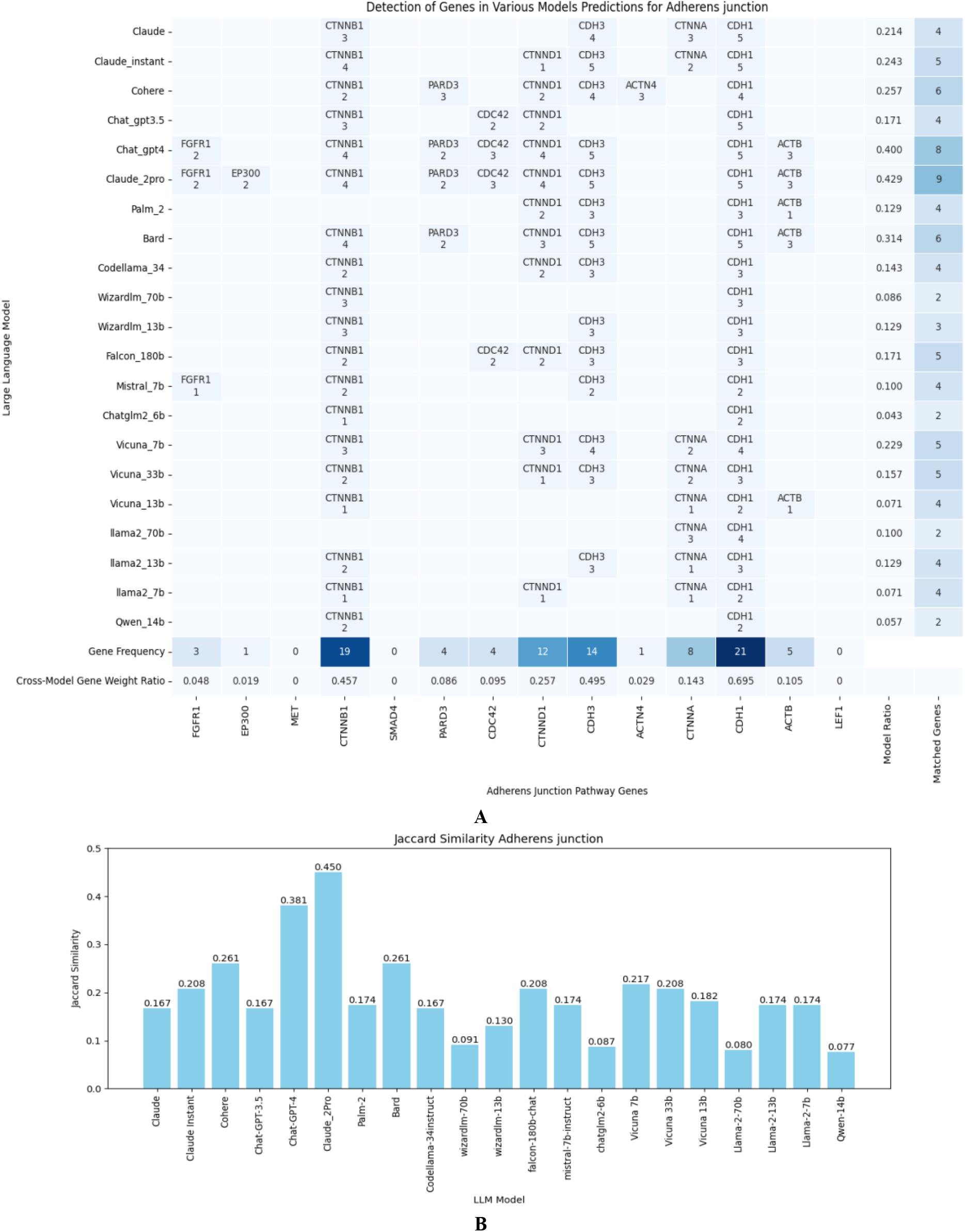
Gene predictions for Adherens Junctions. Part (A) displays prediction accuracy and confidence scores from the 21 API-based and open-source models. The bottom two rows show the frequency of correct gene occurrence among models (one or more correct predictions are counted as success for the model) and among all predictions (cross-model gene weight ratio is the number of correct predictions divided by the number of all predictions, where each model has 5 predictions) for a given gene. The right two columns show the frequency of correct gene occurrence in all predictions (the number of correct predictions divided by the number of all predictions for the model, where each model has 5 predictions), and among genes (one or more correct predictions are counted as success for the gene) for a given model. Part B compares the models through Jaccard similarity scores, assessing their accuracy in matching gene sets specific to Adherens Junctions.

The Jaccard similarity Index, depicted in Figure 2B, offers a comprehensive analysis of the model performance. This index measures the similarity between the predicted genes of each LLM and a reference set of known genes associated with the Adherens junction, thereby providing a statistical basis for comparing their accuracies. All LLMs generated some probable false positives as shown in Table S1, where we examine the unmatched genes to the known genes in the Adherens junction pathway from each model. Overall, the API-based models (the first eight models till Bard) performed better than the open-source models. “Claude_2Pr” and “Chat_gpt4” models emerged as the top performers, predicting more genes correctly and having fewer false positives than other models. The “Claude_2Pr” model topped the list with nine matched genes (FGFR1, PARD3, CDH3, CTNNB1, CDH1, CDC42, EP300, ACTB, and CTNND1) with weights (2, 2, 5, 4, 5, 3, 3, 3, 4), a Jaccard similarity of 0.450, and only six unmatched genes. “Chat_gpt4” matched eight genes (FGFR1, PARD3, CDH3, CTNNB1, CDH1, CDC42, ACTB, and CTNND1) with weights (2, 2, 5, 4, 5, 3, 3, 4), achieving a Jaccard similarity of 0.381 and leaving seven genes unmatched.

In other API-based models, “Claude” identified four genes (CDH1, CTNNA, CDH3, and CTNNB1) with weights (5, 3, 4, 3) and achieved a Jaccard similarity of 0.167, leaving 11 genes unmatched. “Claude Instant” improved slightly, matching five genes (CTNNA, CDH3, CTNNB1, CDH1, and CTNND1) with weights (2, 5, 4, 5, 1), resulting in a Jaccard similarity of 0.208 and 10 unmatched genes. “Coher” outperformed the previous two by identifying six genes (PARD3, CDH3, CTNNB1, CTNND1, CDH1, CTNNA, and ACTN4) with weights (3, 4, 2, 4, 2, 3) and a Jaccard similarity of 0.261, leaving nine genes unmatched. In contrast, “Chat_gpt3.5.” matched four genes (CTNNB1, CDH1, CDC42, and CTNND1) with weights (3, 5, 2, 2), resulting in a Jaccard similarity of 0.167 and 11 unmatched genes.

In the Open-Source category, the standout model was less clear, with several models displaying similar levels of performance. “Codellama_3” matched four genes (CDH3, CTNNB1, CDH1, and CTNND1) with weights (3, 2, 3, 2), yielding a Jaccard similarity of 0.167 and leaving 11 genes unmatched. “Wizardlm_70” identified only two genes (CTNNB1 and CDH1) with equal weights (3, 3), resulting in a lower Jaccard similarity of 0.091 and 13 unmatched genes. “Falcon_180” model matched five genes (CDH3, CTNNB1, CDH1, CDC42, and CTNND1) with weights (3, 2, 3, 2, 2), achieving a Jaccard similarity of 0.208 and leaving 10 genes unmatched. “Mistral_7” matched four genes (FGFR1, CDH3, CTNNB1, and CDH1) with weights (1, 2, 2, 2), resulting in a Jaccard similarity of 0.174 and 10 unmatched genes. The “Vicun” series offered varied results, with “Vicuna_7” matching five genes (CTNNA, CDH3, CTNNB1, CDH1, and CTNND1) with weights (1, 2, 2, 2, 3) and a Jaccard similarity of 0.217.

#### 2.2.2. Tight junction

Figure S1 provides a comprehensive evaluation of 21 LLMs in predicting genes linked to the Tight junction complex, a crucial component in cellular adhesion and signal transduction. “Chat_gpt4” and “Claude_2pr” again emerged as the most proficient, both with a Jaccard similarity of 0.261 and predicting the same subset of genes correctly, closely followed by “Falcon_180” and “Llama2_7”, both with a Jaccard similarity of 0.227. The rest of the models ranged from moderate to low effectiveness in this task. Within the API-based models, “Cohere” achieved a good Jaccard similarity of 0.208. “Claude” and “Claude Instant” had Jaccard similarities of 0.154 and 0.160, respectively. Models like “Palm” and “Bard” lagged, matching fewer genes and thus attaining lower similarity scores. In the open-source category, models like “Codellama_3” and “Wizardlm_70” exhibited moderate performance. The “Mistral_7”, “Chatglm2_6”, and “Vicun” series demonstrated varied outcomes, with “Vicuna_7” and “Vicuna_13” reaching a similarity score of 0.182. “Qwen_14” fell slightly short in this regard. Additionally, Table S2 delved into the unmatched genes for each model.

#### 2.2.3. Gap junction

Figure S2 presents an analysis of 21 LLMs’ capabilities to predict genes associated with the Gap Junction complex, a pivotal element in cellular cohesion and signal transduction. Among these, “Chat_gpt4” and “Claude_2pr” were again identified as the most adept models, with “Llama2_7” showing close competence. Other models’ relative performance was similar to the case of Tight junction. Furthermore, Table S3 complements these insights by examining the unmatched genes.

#### 2.2.4. Cellular Senescence Function

Figure S3 details an extensive evaluation of the same 21 LLMs in predicting genes linked to Cellular Senescence, another critical component in cellular communication and adhesion. “Chat_gpt4” and “Claude_2pr” again led in proficiency, but “Bard” tied their performance. They all have a Jaccard similarity of 0.250. Interestingly, “Chat_gpt4” and “Claude_2pr” predicted the same subset of 6 genes correctly, but “Bard” shared only 5 of them and predicted a different gene correctly, which provided some diversity. The performance of the remaining models varied from moderate to low. Additionally, Table S4 shows the unmatched genes.

#### 2.2.5. Phagosome Function

Figure S4 presents an extensive analysis of 21 LLMs’ abilities to predict genes relevant to the Phagosome Function complex. This complex plays a significant role in cellular adhesion and signal transduction. “Chat_gpt4” stood out for its accuracy, closely followed by “Falcon_180”. After them, “Claude_2pr” and “Llama2_7” tied. Table S5 shows the unmatched genes.

#### 2.2.6. Proteoglycans in Cancer

Figure S5 offers a thorough evaluation of the 21 LLMs concerning their accuracy in predicting genes associated with the Proteoglycans in the Cancer complex, a vital element in cellular adhesion and signal transmission. “Chat_gpt4” and “Claude_2pr” were again the most proficient, followed closely by “Falcon_180” and “Llama2_7”. Table S6 shows the unmatched genes.

#### 2.2.7. Autoimmune Thyroid Disease

Figure S6 provides a comprehensive evaluation of the 21 LLMs in terms of their capacity to predict genes relevant to the Autoimmune Thyroid Disease complex, a crucial aspect of cellular cohesion and communication. ChatGPT-4 and “Claude_2pr” were identified as the most accurate, pursued by “Falcon_180”. In addition, Table S7 shows the unmatched genes.

#### 2.2.8. Overall evaluations

Combining all the results above, we evaluated the overall performance of the twenty-one computational models, dividing them into two distinct categories: API-based and open-source. This detailed assessment aimed to gauge their efficacy in predicting gene regulatory relations and identifying KEGG pathway components, a crucial aspect in the interpretation of complex biological data.

Table 1 shows that the API-based category, ChatGPT-4 emerged as the top performer, demonstrating exceptional prowess in gene regulatory relations with a score of 0.4448, earning it the top rank. It also showed impressive capabilities in KEGG pathway recognition, securing the first rank with a score of 0.2778. Claude-Pro followed closely, excelling particularly in KEGG pathway recognition where it achieved the second-highest score of 0.2657, alongside a strong performance in gene regulatory relations (0.4386, ranking second). Cohere also showed commendable performance, securing the fourth rank in gene regulatory relations (0.3742) and third rank in KEGG pathway recognition (0.2089), highlighting its robust analytical abilities. Other API-based models like Claude-2 and Palm2 displayed moderate performance. Claude-2 ranked third in gene regulatory relations (0.4215) and fifth in KEGG pathway recognition (0.1890). Conversely, Claude instant ranked fifth in gene regulatory relations (0.3263) and sixth in KEGG pathway recognition (0.1519). Palm2 and Bard, showing moderate efficacy, ranked sixth and seventh in gene regulatory relations, respectively, but fared slightly better in KEGG pathway recognition, placing eight and fourth.

**Table 1:** Comparative performance of twenty-one computational models in predicting gene regulatory relations and recognizing KEGG pathways, Categorized as API-based and open-source models.

| Type | Model | Gene Regulatory Relations (F1) | Gene Regulatory Relations (Rank) | KEGG Pathway Recognition String (Mean) | KEGG Pathway Recognition (String) Rank |
| --- | --- | --- | --- | --- | --- |
| API base | Claude-2 | 0.4215 | 3 | 0.1890 | 5 |
|  | Claude-instant | 0.3263 | 5 | 0.1519 | 6 |
|  | Chat_GPT3.5 | 0.1485 | 8 | 0.1358 | 7 |
|  | Cohere | 0.3742 | 4 | 0.2089 | 3 |
|  | **ChatGPT-4** | **0.4448** | **1** | **0.2778** | **1** |
|  | Claude-Pro | 0.4386 | 2 | 0.2657 | 2 |
|  | Palm2 | 0.2614 | 7 | 0.1048 | 8 |
|  | Bard | 0.2785 | 6 | 0.1923 | 4 |
| Open source | Codellama-34 | 0.1435 | 6 | 0.1622 | 5 |
|  | Wizardlm-70b | 0.0449 | 13 | 0.1345 | 8 |
|  | Wizardlm-13b | 0.0962 | 11 | 0.1459 | 7 |
|  | **Falcon-180b** | **0.2787** | **1** | **0.2237** | **1** |
|  | Mistral-7b | 0.1226 | 8 | 0.1428 | 9 |
|  | chatglm2-6b | 0.1821 | 3 | 0.0881 | 12 |
|  | Vicuna-7b | 0.1225 | 9 | 0.1744 | 4 |
|  | Vicuna-33b | 0.1405 | 7 | 0.1658 | 6 |
|  | Vicuna-13b | 0.1806 | 4 | 0.1540 | 10 |
|  | llama-2-70b | 0.1583 | 5 | 0.1540 | 11 |
|  | llama-2-13b | 0.0912 | 12 | 0.1712 | 3 |
|  | llama-2-7b | 0.1923 | 2 | 0.2207 | 2 |
|  | Qwen-14b | 0.1013 | 10 | 0.0842 | 13 |

In open-source models, Falcon-180b led the pack, outperforming its counterparts in both gene regulatory relations (0.2787, rank 1) and KEGG pathway recognition (0.2237, rank 1). This underscores its potential as a comprehensive analytical tool in genomic studies. Llama2-7b also displayed a strong performance, ranking second in both gene regulatory relations (0.1923) and KEGG pathway recognition (0.2207). Codellama-34 and Chatglm2-6b also demonstrated robust performances in gene regulatory relations, scoring 0.1821 and 0.1435, respectively. The Vicuna and Llama2 series exhibited varying levels of performance. Notably, Vicuna-7b, despite ranking nine in gene regulatory relations, showed a specific strength in KEGG pathway recognition, ranking fourth. Qwen-14b, ranking the lowest in both gene regulatory relations (0.0100) and KEGG pathway recognition (0.0842), highlighted some of the challenges faced by models in interpreting complex genomic data.

Overall, this evaluation sheds light on the diverse capabilities of the computational models, with certain models like ChatGPT4 and Claude-Pro showcasing high accuracy, and others such as Qwen-14b exhibiting lower effectiveness. The variation in gene prediction confidence across different models underscores the importance of selecting the right computational tools for specific tasks in genomic research, guiding the understanding of gene regulation and pathway analysis complexities in high-impact scientific studies.

## 3. DISCUSSION

In biomedical research, LLMs have been explored in extracting and analyzing complex biological data. Researchers often use LLMs, especially ChatGPT, for their biomedical research problems. However, to our knowledge, no systematic assessment has been conducted for applying LLMs in biomedical knowledge retrieval. This study has comprehensively evaluated the efficacy of 21 LLMs, both API-based and open-source, in predicting gene regulatory relations and KEGG pathway components. A critical aspect of our research is the use of the ground truth in the objective benchmarking. By employing metrics like accuracy, precision, recall, F1 score, and the Jaccard similarity index, we have provided benchmarks and quantifiable comparisons of these models against established ground truths. A notable finding from this study is the disparity in performance among different models. API-based models like Claude-Pro and ChatGPT4 have demonstrated exceptional capabilities, potentially thanks to their large model sizes and inclusion of training text specific to biomedical research [17]. It is also noted that ChatGPT-4 and Claude-Pro have many similar predictions (Figures 1, S1-S6), probably because they shared similar methods and training data. In contrast, although other models performed worse, they may offer complementary correct predictions, indicating a potential to integrate several LLMs to achieve better meta-analysis results.

While LLMs offer significant potential in biomedical research, their deployment must be carefully selected and managed, focusing on enhancing accuracy and mitigating biases. The performance disparity among the evaluated models, particularly between the leading API-based models like Claude-Pro and ChatGPT4, and the best-performing open-source models Falcon-180b and Llama2-7b, underscores the significance of model selection tailored to the specific demands of biomedical tasks. Our study also identified limitations of LLMs that need to be addressed. The accuracy and specificity of these models, particularly in identifying complex biomedical terms, require further improvement. This could potentially be achieved through more specialized training on targeted datasets [22]. Integrating LLMs with existing bioinformatics tools and databases could lead to a more comprehensive approach to gene function and pathway analysis [23]. Additionally, the continual evolution of LLM algorithms and their adaptation to specific biomedical applications will be crucial to keep pace with the rapid advancements in the field [24].

There are some limitations of this study as well. The study’s focus on a specific set of 200 genes and 7 pathways presents a limitation in terms of the breadth of the research. While this selection allows for in-depth analysis of these genes, it may not fully capture the diversity and complexity inherent in biomedical research at large. The findings derived from this constrained set might not be entirely applicable or representative of other gene sets, which could exhibit different patterns or interactions. This limitation points to the need for caution in generalizing the study’s results to the broader field of biomedicine and beyond. The use of different prompts in the study introduces a variable that could affect the consistency and reliability of the results. Since LLMs’ responses can vary significantly based on the input prompts, the diversity in the prompts used might lead to varied responses, making it challenging to accurately assess the overall performance and capabilities of the LLMs under different prompts. This variability could potentially skew the study’s findings, as the LLMs might perform better or worse depending on the specific prompts used.

In summary, our study underscores the great potential of LLMs in biomedical knowledge mining from the literature, including gene regulatory network analysis and pathway mapping. The integration of LLMs into the biomedical research ecosystem is poised to transform the way complex biological data is analyzed and interpreted, thereby accelerating scientific discoveries and innovations in healthcare. This necessitates continuous evolution and refinement of LLM algorithms, ensuring their relevance and efficacy in the rapidly advancing field of biomedical research.

## 4. MATERIALS AND METHODS

We developed a pipeline to evaluate 21 LLMs, as shown in Figure 3. In this study, we extensively utilized the KEGG Pathway Database as the primary source for establishing the ground truth. We first analyzed gene regulatory relationships, with an emphasis on processes such as activation, inhibition, and phosphorylation. To this end, we established interactions from the KEGG pathway. Then we selected seven crucial biomedical terms within the KEGG pathway. Prompts were crafted and applied to each LLM to extract pertinent data. We then comprehensively evaluate the performance of these models using statistical measures such as accuracy, precision, recall, and F1 scores, along with the Jaccard similarity index for matching genes.

**Figure 3:**
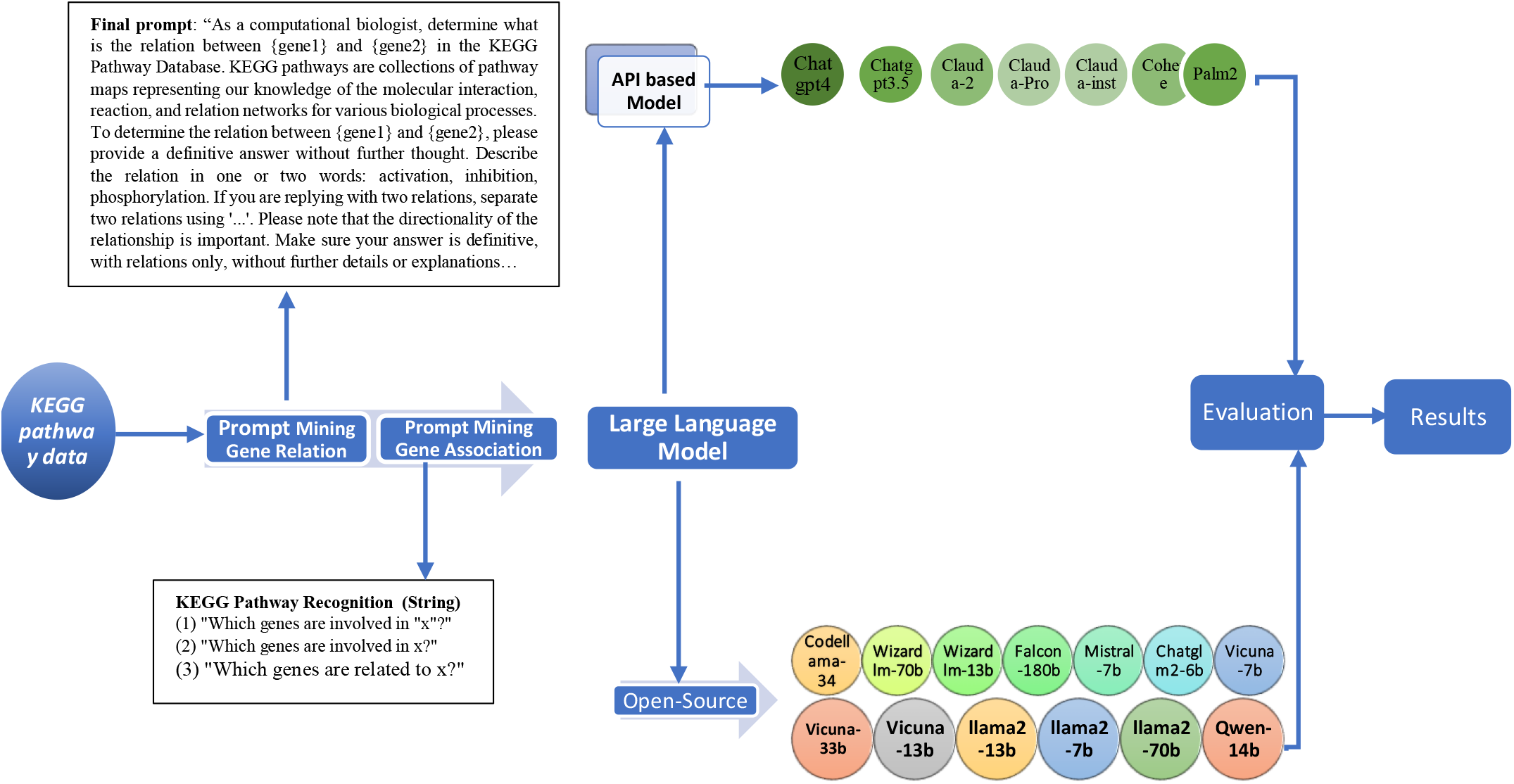
An assessment workflow for LLMs to analyze biomedical data. The process starts with prompts to LLMs for gene relationship prediction and recognition of genes associated with biomedical terms. The LLMs used in this study include both API-based and open-source models. The results are evaluated comprehensively and rigorously.

### 4.1. Evaluating Gene Regulatory Relations

We employed the KEGG Markup Language (KGML) for effective data extraction and integration. A key element of our methodology was the selection of 200 gene triplets from diverse pathway maps, representing various relationships. These triplets were meticulously chosen to exemplify three principal types of gene interactions: activation, inhibition, and phosphorylation, thereby providing a comprehensive framework for our experiments. The selection’s relevance and accuracy were validated through their documented presence in KEGG pathway diagrams, as discussed in our previous paper’s [26]. This data includes 200 gene triplets, covering activations, inhibitions, and phosphorylations (with 80 instances each for activations and inhibitions, and 40 for phosphorylations). The complete list of 200 gene triplets can be found in Table S8.

Effective prompt writing is crucial in large language models, as a well-constructed prompt can extract more valuable information on gene regulatory relations. We adopted a focused approach in prompt construction, emphasizing the role of precise and concise questioning. This method is particularly evident in our use of the Few-shot and Role Prompting techniques, which aim to extract highly specific information from LLMs. An example of this approach is the prompt we designed for a computational biologist: “Determine the relationship between {gene1} and {gene2} in the KEGG Pathway Database.” The KEGG pathways encompass a variety of biological processes, detailing molecular interactions, reactions, and networks. The prompt specifically requests a straightforward categorization of the relationship between two genes, using terms like ‘activation,’ ‘inhibition,’ or ‘phosphorylation,’ and emphasizes the significance of the relationship’s directionality. This method ensures that responses are clear and limited to essential information without additional detail or explanation. The prompt is not optimized for any LLM. Based on some initial trials, the final prompt is as follows:

#### Final prompt

“As a computational biologist, determine what is the relation between {gene1} and {gene2} in the KEGG Pathway Database. KEGG pathways are collections of pathway maps representing our knowledge of the molecular interaction, reaction, and relation networks for various biological processes. To determine the relation between {gene1} and {gene2}, please provide a definitive answer without further thought. Describe the relation in one or two words: activation, inhibition, phosphorylation. If you are replying with two relations, separate two relations using ‘…’. Please note that the directionality of the relationship is important. Make sure your answer is definitive, with relations only, without further details or explanations. Make sure your answer is definitive, composed of ‘activation’, ‘inhibition’, ‘phosphorylation’ or ‘no information’ without further details or explanation.”

#### Question

Q: What effect does gene CCR5 have on gene GNB3?

Q: What effect does gene LIMK1 have on gene CASP1?

Q: What effect does gene IFNB1 have on gene IL10RA?

Q: What effect does Gene IFNA4 have on gene IL10RA?

### 4.2. Evaluations KEGG Pathway Recognition

We focused on identifying seven key biomedical terms within the KEGG Pathway framework. These terms are Adherens Junction, Tight Junction, GAP Junction, Cellular Senescence, Phagosome, Proteoglycans in Cancer, and Autoimmune Thyroid Disease. The comprehensive ground truth data for all the examined biomedical terms is detailed in Figures 1 and S1-S6. We employed 21 different LLMs to assess their accuracy in identifying these specific biomedical terms. Our evaluation was based on a detailed comparison with verified data related to these terms. Our analysis included not only instances where the models successfully identified the correct gene associations but also cases where they failed to match genes. To improve the reliability of our assessments, we repeated one query five times, assigning a weighted significance to each gene based on its occurrence. This process involved querying different biomedical terms five times each. The prompt used is as follows:

“Which genes are involved in x?”

The queries were as follows:

- “Which genes are involved in ‘Adherens junction’?”
- “Which genes are involved in ‘Tight junction’?”
- “Which genes are involved in ‘GAP junction’?”
- “Which genes are involved in ‘Cellular senescence’?”
- “Which genes are involved in ‘Phagosome’?”
- “Which genes are involved in ‘Proteoglycans in cancer’?”
- “Which genes are involved in ‘Autoimmune thyroid disease’?”

### 4.3. Large Language Models

We investigated the capabilities of advanced LLMs, focusing on both API-based and open-source models, including smaller, specialized models tailored for the biomedical domain. Our goal was to assess their proficiency in various biological tasks, with a particular emphasis on pathway knowledge and gene regulatory relationships [27]. The technical specifications of these models are detailed in Table 2.

**Table 2:**
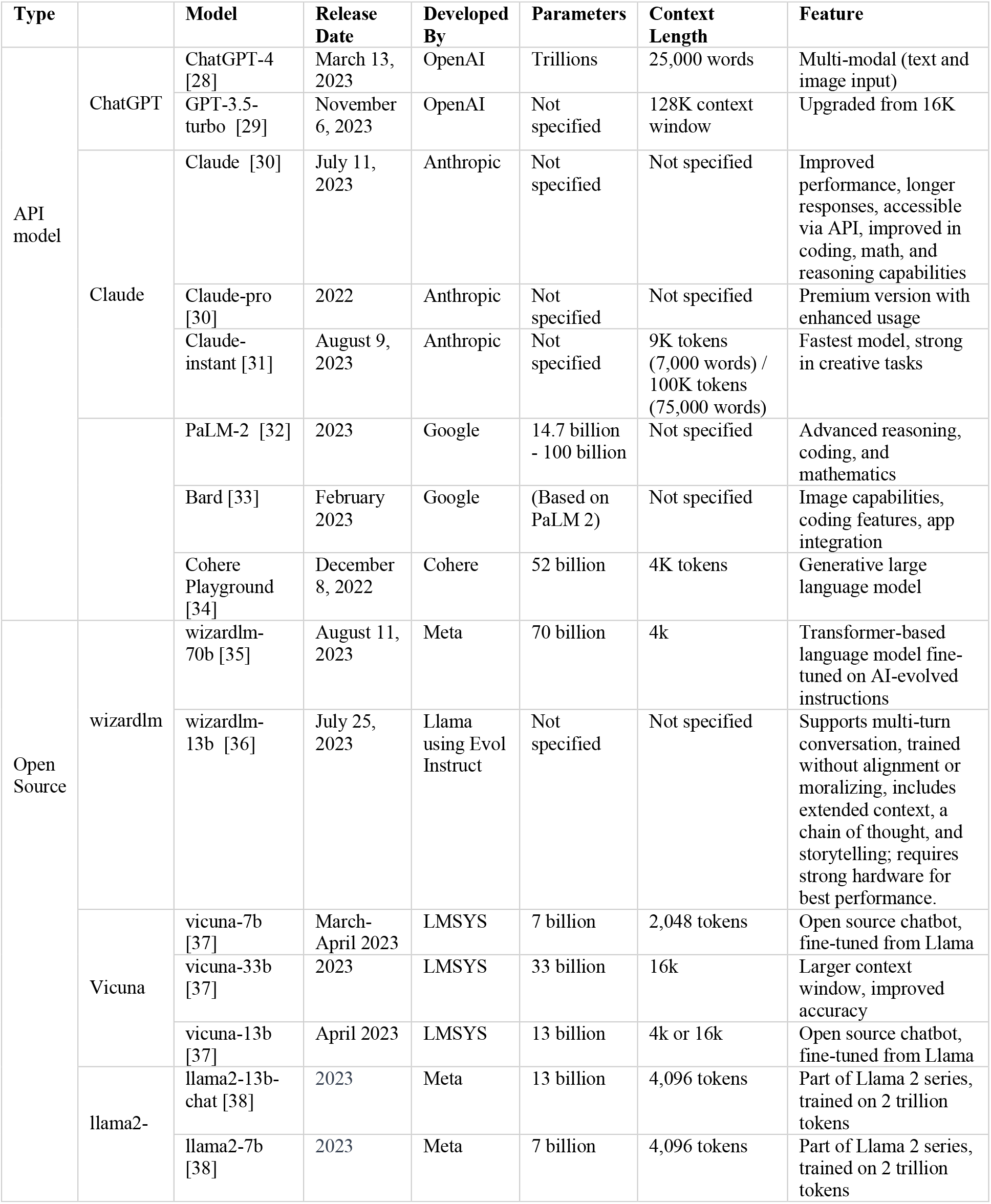
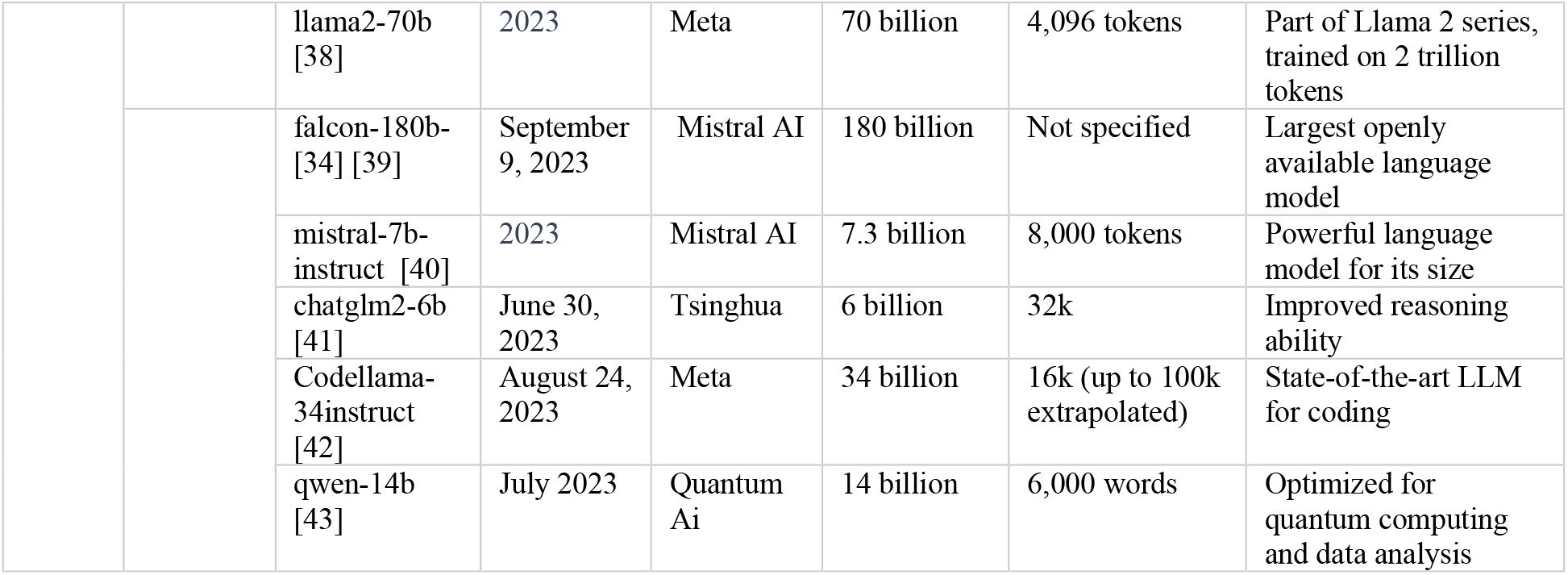
Overview of Leading Language Models Specifications and Unique Features.

### 4.4. Evaluations

In our analysis, we assessed the performance of various LLMs in predicting gene regulatory relationships within KEGG pathways [44]. The evaluation metrics were calculated by comparing the ground truth data (ground truth), which represents the actual gene relationships, with the predictions (predicted labels) made by each LLM. Here’s how each metric was contextualized:

#### Precision

Precision in our study quantifies the accuracy of the LLMs in correctly predicting gene regulatory relationships. It reflects the model’s ability to minimize false positives, ensuring that the relationships it predicts are indeed relevant and accurate:

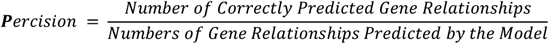

#### Recall (Sensitivity)

Recall measures the LLMs’ ability to identify all existing gene regulatory relationships in the KEGG pathways. It is crucial for ensuring that the model captures as many true regulatory relationships as possible, minimizing missed detections (false negatives):

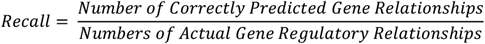

#### F1 Score

The F1 Score balances the trade-off between Precision and Recall. It provides a single metric that encapsulates the model’s overall performance in accurately identifying gene regulatory relationships while minimizing both false positives and false negatives:

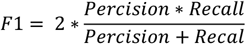

These metrics were specifically chosen and calculated for each LLM to evaluate their effectiveness in the nuanced task of predicting gene regulatory relationships within KEGG pathways.

#### Jaccard similarity

We employed the Jaccard similarity metric to evaluate the performance of LLMs in recognizing strings related to KEGG Pathways [45]. We specifically focus on the seven biomedical terms. The central objective of this evaluation was to analyze the gene overlaps, by comparing the genes predicted by various models to the actual, known genes - a set we refer to as the “ground truth”. The Jaccard similarity measures the proportion of common genes to the total unique genes identified by both the model predictions and the actual data:

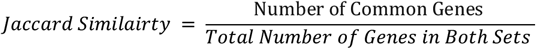

## AUTHOR CONTRIBUTIONS

All authors have accepted responsibility for the entire content of this manuscript and approved its submission.

## ACKNOWLEDGEMENTS

This work was supported by the National Institute of Health (R01-LM013392, R35-GM126985 and P30DK092950).

## CONFLICT OF INTEREST STATEMENT

Authors state no conflict of interest.

## DATA AVAILABILITY STATEMENT

The data structure concerning gene pathways and biomedical terms, as developed and analyzed in our study, can be found on the KEGG website and in the Supplemental File.

## ETHICS STATEMENT

This article does not contain any studies with human or animal materials performed by any of the authors.

## Notes

### Competing Interest Statement

The authors have declared no competing interest.

